# Vib2Sound: Separation of Multimodal Sound Sources

**DOI:** 10.1101/2025.05.08.652866

**Authors:** Mai Akahoshi, Yuhang Wang, Longbiao Cheng, Anja T. Zai, Richard H. R. Hahnloser

## Abstract

Understanding animal social behaviors, including vocal communication, requires longitudinal observation of interacting individuals. However, isolating individual-level vocalizations in complex environments is challenging due to background noise and frequent overlaps of coincident signals from multiple vocalizers. A promising solution lies in multimodal recordings that combine traditional microphones with animal-borne sensors, such as accelerometers and directional microphones. These sensors, however, are constrained by strict limits on weight, size, and power consumption and often lead to noisy or unstable signals. In this work, we introduce a neural network-based system for sound source separation which leverages multi-channel microphone recordings and body-mounted accelerometer signals. Using a dataset of zebra finches recorded in a social setting, we demonstrate that contact sensing largely outperforms conventional microphone-array recordings. By enabling the separation of overlapping vocalizations, our approach offers a valuable tool for studying animal communication in complex naturalistic environments.

## 1. INTRODUCTION

Understanding social behaviors in animals requires longitudinal observation of interactions in group settings. Among the behaviors of interest, vocal communication has attracted widespread interest across disciplines [1, 2]. Traditionally, vocal behaviors are recorded using microphones that capture all sounds generated in the scene. However, in social settings where multiple animals can vocalize simultaneously, such recordings often contain overlapping vocalizations and environmental background noise, making it difficult to attribute vocalizations to specific individuals.

Microphone array-based approaches have been proposed to meet this challenge, as microphone arrays capture sound at multiple spatial locations, allowing the exploitation of time and amplitude differences between microphones to estimate the direction or position of a vocal source [3]. While microphone arrays combined with neural networks have shown superior performance in human speech separation [4, 5], their application in bioacoustics is relatively limited. Overlapping vocalizations among individuals of the same species are especially hard to separate due to their high acoustic similarity, forcing researchers to exclude such overlapping vocalizations from analysis [6, 7]. To address this issue, body-mounted sensors, such as miniature microphones and accelerometers, have been proposed for use in rats [8], mice [9], birds [10], and humans [11]. Although such sensors can provide valuable individual-level data, they are often compromised in signal quality compared to microphone signals due to strict constraints on weight, size, and power consumption. Also, such sensors often signal low-frequency body vibrations or near-field sound signals that differ from traditional far-field microphone signals, limiting the comparability between studies. Therefore, an integrative approach is required that takes advantage of the complementary strengths of the microphone and body-mounted sensor data to extract clean vocalizations of each individual.

This challenge closely parallels Target Speech Extraction (TSE) in human speech processing. TSE aims to isolate the speech of a target speaker from a mixture of multiple speakers under noise and reverberation. Recent TSE systems exploit multiple cues to identify and separate the speakers in the mixture [12], including audio cues from pre-recorded utterances of the target speaker [13, 14], spatial cues from microphone arrays [15, 16] or from video recordings of the overall scene [17], and visual cues such as lip movements captured through video analysis [18, 19]. Neural networks (NNs) typically integrate these diverse cues into robust pipelines for speech signal separation.

Following the TSE frameworks, we propose a vocalization separation system that uses body-mounted sensor signals as cues to separate overlapping vocalizations recorded with high-quality wall-mounted microphones. In this approach, accelerometer-based sensors mounted on animals record vocalizations via body vibrations, providing individual-specific cues. Additionally, multiple microphones provide spatially distinct audio recordings, potentially aiding in individual discrimination and sound separation. Building on an existing TSE framework [20], we propose Vib2Sound, a neural network for vocalization separation specifically designed to integrate individual-specific accelerometer cues and spatial information from microphone arrays.

We evaluate the efficacy of Vib2Sound on data collected in BirdPark [10], a sound-isolated environment and system for synchronized multimodal recordings from animal groups. Each zebra finch in this setup carries an accelerometer mounted on its body, capturing the low-frequency vibration associated with vocalizations (Fig. 1). Prior analysis of BirdPark data showed that microphones provide more reliable signals of vocalizations than accelerometers, questioning the usefulness of contact sensing for vocal analysis.

**Fig. 1.**
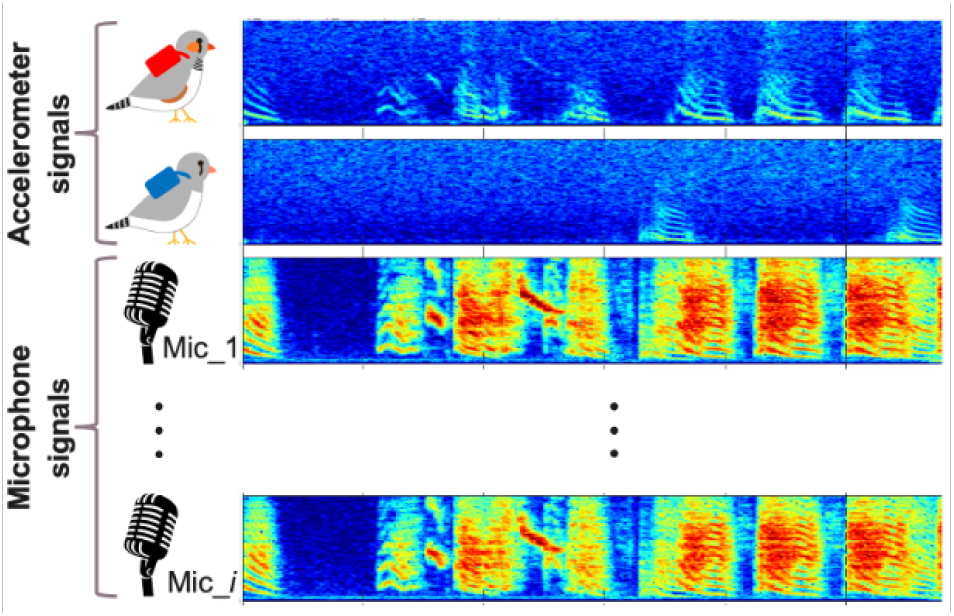
Example synchronized data stream recorded in BirdPark [10], showing two overlapping vocalizations from different individuals. The top two rows show spectrograms of body-mounted accelerometer signals, while the bottom spectrograms are derived from wall-mounted microphones.

Specifically, our objectives are (1) to evaluate the performance of Vib2Sound in accurately separating overlapping vocalizations, (2) to investigate the extent to which body-mounted accelerometer cues enhance vocalization attribution compared to microphone-only methods, and (3) to assess the generalizability of Vib2Sound to realistic scenarios encountered in naturalistic settings. By addressing these objectives, our study aims to quantify the usefulness of contact sensing and to establish a robust methodological framework to advance individual-level analyses of vocal communication in animal groups, contributing valuable insights to research in behavioral ecology and bioacoustics.

## 2. METHODS

### 2.1. Proposed model

The structure of Vib2Sound^1^ is summarized in Fig. 2. Vib2Sound is based on VoiceFilter [20], but replaces the speaker-specific vocal embeddings with accelerometer-derived vibration cues and rearchitects the model to operate on multi-channel audio. As inputs, Vib2Sound takes the mixed multi-channel microphone signals (Mic) and the respective accelerometer signals (Acc) from two birds. The network outputs two time-frequency masks that we used to predict the separated vocalizations of the two birds when applied to the mixture spectrogram. The network is trained to minimize the mean squared error (MSE) between the predicted spectrograms and the two ground-truth spectrograms. We applied an inverse Short-Time Fourier Transform (iSTFT) to generate demixed audio waveforms.

**Fig. 2.**
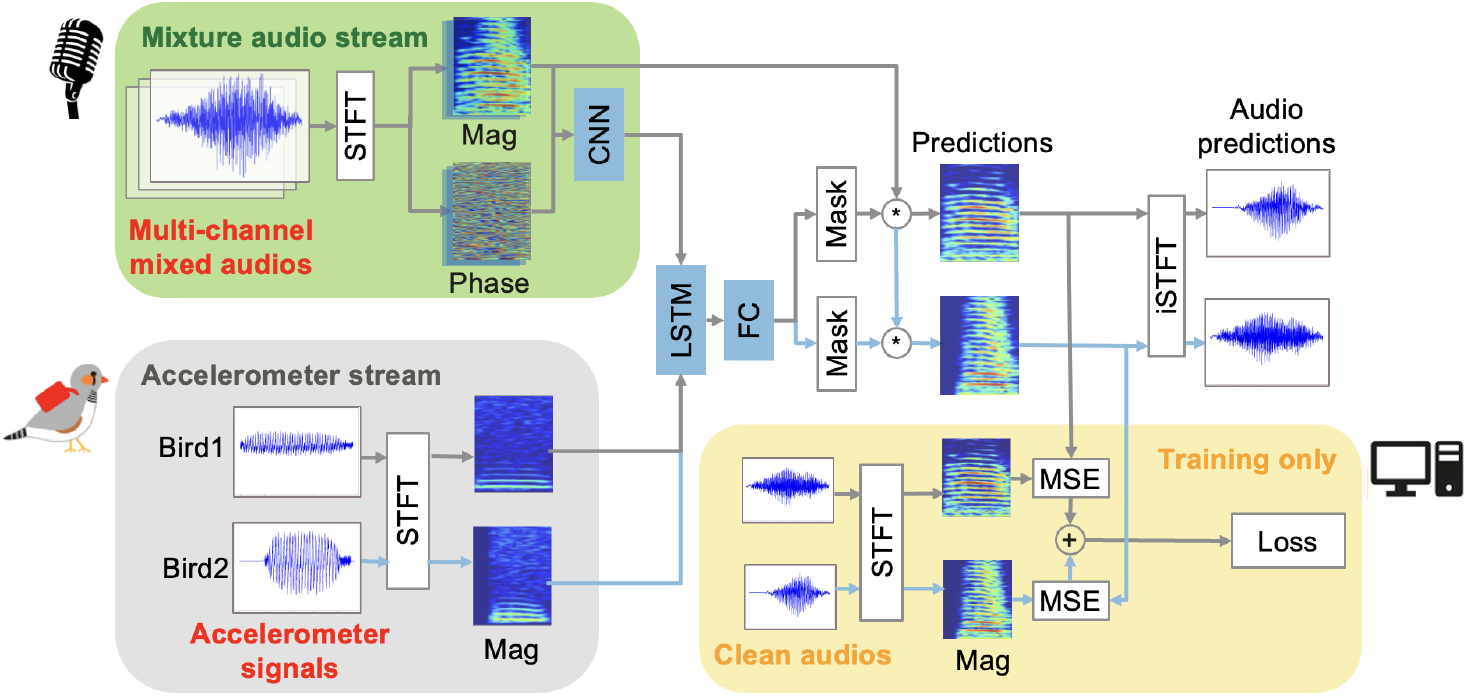
Structure of Vib2Sound. The CNN output of the mixture audio stream and the accelerometer spectrograms all feed into the LSTM from where two masks are computed that we used to generate a predicted source spectrogram for each bird (gray and light blue arrows). The MSE losses of these predictions are summed and used as the training loss to be minimized. Red font indicates inputs to the model and orange shading indicates the loss computation during training.

The system consists of an 8-layer Convolutional Neural Network (CNN), 1 Long Short-Term Memory (LSTM) layer, and a 2-layer Fully Connected (FC) block. It computes the magnitude (Mag) and phase spectrograms of the mixture signals from all channels and passes these to the CNN to extract acoustic features (mixture audio stream). It concatenates the magnitude spectrograms of the accelerometer signals (accelerometer stream) to the CNN output and feeds it to the LSTM layer. The output of the LSTM layer feeds into the FC block that generates soft masks. The soft mask outputs are then applied to the magnitude mixture spectrogram from the first microphone channel to predict the source spectrograms. We turned the predicted source spectrograms into audio waveforms using the phase signal of the mixture spectrogram.

### 2.2. Dataset

We constructed a training dataset from clean isolated vocalizations. Following VoiceFilter, we artificially mixed microphone signals by mixing pairs of randomly chosen clean vocalizations of two different birds, Fig. 3. To simulate realistic timing variability, we generated training samples by positioning the shorter audio segment at a random onset within the time span of the longer one.

**Fig. 3.**
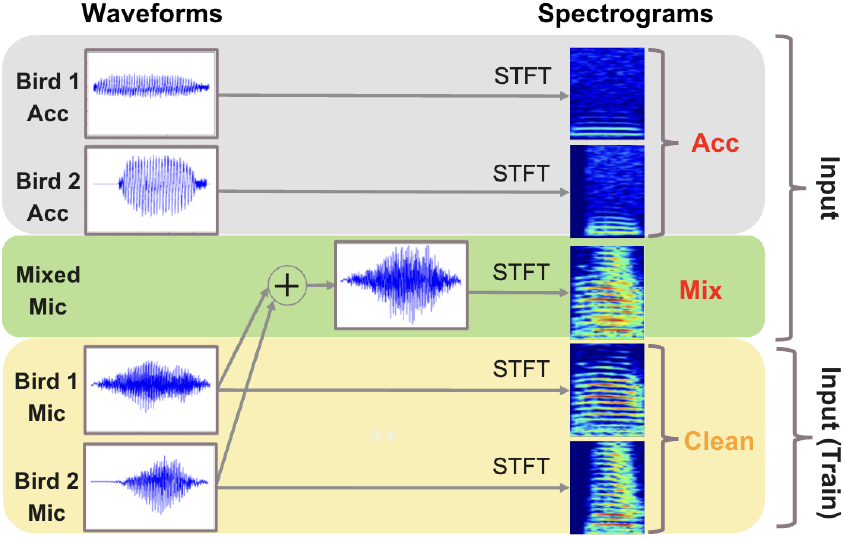
We generated artificial sound mixtures (Mix) for training by aligning vocalizations from pairs of birds and summing their microphone signals.

Clean audios were extracted from recordings of 10 mixed-sex bird pairs in BirdPark, collected over 4 to 13 days per pair, with a sampling rate of 24414 Hz. We detected vocalizations using WhisperSeg [21], an automated vocal activity detection system. We manually selected non-overlapping clean vocalizations and assigned them to training, validation, and evaluation datasets. For each bird, we reserved 10 % of the data as the evaluation dataset and 4.5 % as the validation dataset. We divided each dataset into two subsets based on their microphone configurations, since spatial cues are highly sensitive to microphone position.

Dataset1 (14 birds): This setup includes 4 microphones. Mic1 and mic2 were centered on top of the right/front walls of the arena, mic3 on the back wall of the surrounding isolation box, and mic4 was installed on the ceiling of the box. The number of vocalizations per bird ranged from 430 to 1582 in the training dataset, and in total there were 8894 vocalizations in the training, 462 in the validation, and 1033 in the evaluation dataset.

Dataset2 (6 birds): Mic1-4 are positioned above the 4 walls of the arena and mic5 on the ceiling of the isolation box. The number of vocalizations per bird ranged from 454 to 3263 in the training dataset, and in total there were 8046 vocalizations in the training, 420 in the validation, and 938 in the evaluation dataset.

### 2.3. Training

We use the standard Short-Time Fourier Transform (STFT) with length of 384 samples and a hop size of 96 samples to convert both the multi-channel microphone signals and the accelerometer signals into complex (magnitude and phase) time–frequency representations. We applied a 200 Hz high-pass filter to the accelerometer signal to remove low-frequency noise arising from radio signal transmission.

We trained one model for each of the two datasets (see Section 2.2). The batch size used for training was set to 4. We used the Adam optimizer with a learning rate of 0.001 and trained the models for 200,000 steps.

### 2.4. Evaluation

#### 2.4.1. Distance metrics

To evaluate the performance associated with diverse training strategies and models (see Section 2.4.2), we computed the distance score between the clean audio magnitude spectrograms and the predicted magnitude spectrograms on a time-bin basis. For each time bin *t*, we calculated the distance between the clean spectrum **S**_clean_(*t*) and the predicted spectrum **S**_pred_(*t*), and then we averaged the distances across all time bins of the vocalization:

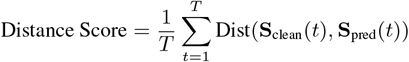

where *T* is the total number of time bins spanned by the vocalization, and Dist denotes the chosen distance metric (Euclidean, Cosine, or Spearman distance).

To assess performance, each model was evaluated on 1,000 artificial mixtures from the corresponding evaluation dataset. Each model generated 2,000 predicted vocalizations (two vocalizations from different birds per mixture), on which we evaluated distance scores.

#### 2.4.2. Baseline models

We compared our proposed approach with the following ablations and baselines.

Single-channel Vib2Sound (ablation): To study the contribution of multi-channel microphone signals to the model, we constructed a single-channel Vib2Sound by feeding only the magnitude spectrogram from the first microphone into the CNN, which was then used for both training and inference.

Audio-only Vib2Sound (ablation): Similarly, we ablated the accelerometer stream to evaluate the contribution of accelerometer signals. In this model, 2 LSTMs and FCs receive CNN output in the mixture audio stream, and each of them produced a mask. We used permutation invariant training (PIT) to ensure correct loss computation. We trained and evaluated both multi-channel and single-channel versions of audio-only model.

TF-GridNet [22]: TF-GridNet is a state-of-the-art speech separation model that leverages microphone array inputs. TF-GridNet operates in the time-frequency domain and involves two multi-channel-input single-channel-output deep neural networks sandwiching a beamformer.

### 2.5. Real-world mixtures

In addition to testing the model on artificially mixed vocalizations, we assessed its performance on 196 naturally overlapping calls extracted from our recordings. Since ground-truth clean sources are not available in such cases, we assessed the plausibility of the separated outputs by analyzing vocal pitch, a widely used metric in birdsong research [23, 24]. We focused on a harmonic call type and extracted 196 real mixtures containing this call from a single bird. To evaluate the Vib2Sound’s separation quality, we compared the pitch distribution of the separated microphone signals with the pitch distribution derived from the microphone signal of 290 clean instances of the same vocalization type from the same bird. This allowed us to test whether Vib2Sound captures subtle pitch statistics and thereby enhances the signal quality relative to the original overlapping signals. Pitch was extracted using Parselmouth [25], a Python interface to Praat [26], a widely used tool for birdsong analysis.

## 3. RESULTS AND DISCUSSION

### 3.1. Performance on artificial data

Fig. 4 shows a representative vocal mixture on which Vib2Sound successfully separates the overlapping vocalizations from two individuals. The predicted spectrograms closely match the ground-truth spectrograms with minimal residuals, indicating accurate reconstruction.

**Fig. 4.**
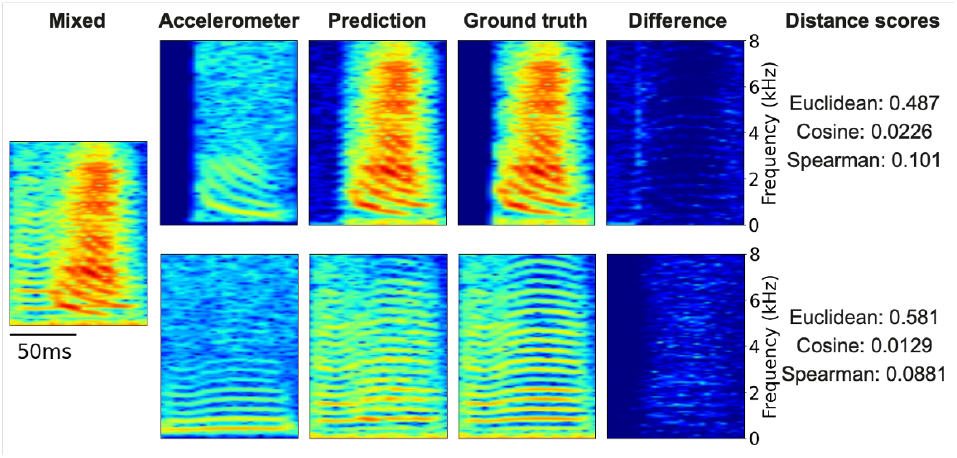
Example of Vib2Sound’s separation of an artificial mixture. From left to right, the spectrograms show: the artificially mixed microphone input, associated accelerometer signals, predicted vocalizations, ground-truth vocalizations, and absolute differences between the predicted and ground-truth spectrograms.

We summarize the performance results on artificial mixtures, including distance scores, Multiply-Accumulate operations (MACs), and parameter sizes, in Table 1. The table also includes distance scores between the ground-truth and the unseparated mixture as a reference (row 1). The results indicate that multi-channel and single-channel Vib2Sound perform nearly identically (row 2 vs. 3). However, when the accelerometer signal is removed, the benefit of multi-channel audio input becomes evident (row 4 vs. 5), suggesting that the model primarily relies on the single microphone channel together with accelerometer input when available.

**Table 1.** Performance comparison of Vib2Sound, ablations and baselines. Bold values indicate the best performance. MACs and parameter sizes are computed for the 5-channel microphone inputs of 200 ms.

|  | dataset1 |  |  | dataset2 |  |  | MACs (G) | params (M) |
| --- | --- | --- | --- | --- | --- | --- | --- | --- |
|  | Euclidean | Cosine | Spearman | Euclidean | Cosine | Spearman |  |  |
| Mixture | 1.665 | 0.0450 | 0.212 | 1.843 | 0.0574 | 0.215 | - | - |
| multi-channel Vib2Sound | **0.637** | **0.0249** | **0.146** | **0.527** | **0.0228** | **0.125** | 1.54 | 7.99 |
| single-channel Vib2Sound | 0.642 | **0.0249** | 0.147 | 0.534 | 0.0229 | 0.126 | 1.53 | 7.98 |
| audio-only multi-channel Vib2Sound | 1.237 | 0.0396 | 0.189 | 1.197 | 0.0470 | 0.182 | 1.52 | 14.19 |
| audio-only single-channel Vib2Sound | 1.377 | 0.0417 | 0.196 | 1.502 | 0.0522 | 0.192 | 1.52 | 14.19 |
| TF-GridNet (200000 steps) | 1.269 | 0.0512 | 0.269 | 1.238 | 0.0564 | 0.236 | 38.68 | 8.32 |
| TF-GridNet (800000 steps) | 1.050 | 0.0415 | 0.221 | 1.032 | 0.0470 | 0.209 | 38.68 | 8.32 |

The results further show that Vib2Sound largely outperforms TF-GridNet across all distance metrics while being computationally more efficient. As shown in rows 6 and 7, TF-GridNet required substantially more training steps for its loss to converge. Although longer training improved TF-GridNet’s results, visual inspection revealed that it captured the vocalization boundaries but showed limitations in reconstructing fine-grained acoustic structure. In contrast, Vib2Sound successfully reconstructed detailed vocalization patterns by leveraging accelerometer signals.

### 3.2. Performance on real-world mixture data

Figure 5 shows the pitch distributions for clean calls, overlapping calls, and calls separated by Vib2Sound from these overlapping calls. The reconstructed calls exhibit pitch statistics closely resembling those of clean calls. Kolmogorov-Smirnov tests confirmed that predicted calls are significantly more similar to clean calls than overlapping calls are to clean calls (p = 0.283 vs. p *<* 0.001, respectively). This demonstrates that Vib2Sound can effectively process real-world overlapping vocalizations, producing separated signals suitable for downstream analysis.

**Fig. 5.**
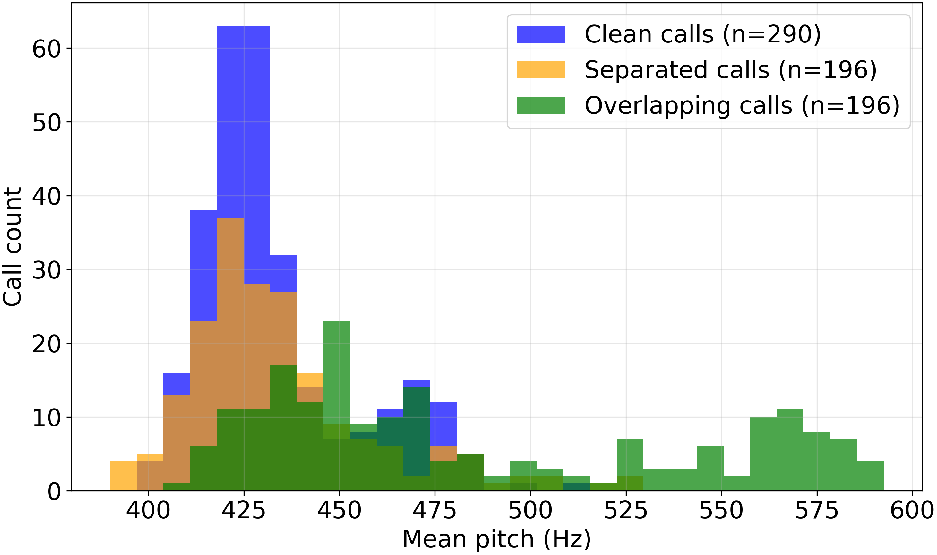
Mean pitch distribution of clean calls, overlapping calls and calls separated by Vib2Sound.

## 4. CONCLUSION

In this work, we proposed Vib2Sound, a sound source separation model for animal vocalizations that works on microphone signals and leverages signals from body-mounted sensors as guiding cues. Evaluated on artificial mixtures of vocalizations of different individuals, our model outperformed a state-of-the-art speech separation model that relies on microphone-array inputs. Furthermore, we demonstrated the reliability of our approach on real-world overlapping vocalizations.

In conclusion, our work highlights the effectiveness of body-mounted sensor signals for reconstructing individual vocalizations from recorded mixtures, which can constitute more than 40% of cases in groups of 8 birds [10]. Vib2Sound showcases the benefit of vibration sensing, even though prior work found animal-borne signals less reliable for detecting vocalizations than remote microphone signals [10]. But since missed vocalizations on accelerometer channels constitute less than 5% of cases [10], the usefulness of Vib2Sound for processing the remaining 95% of detected vocalizations seems obvious. Our work is therefore expected to facilitate the analysis of vocal communication in social animals, promoting deeper understanding of their social interactions and behaviors in natural settings.

## Footnotes

1 Code and scripts are available at https://gitlab.switch.ch/hahnloser-songbird/birdpark/vib2sound

